# RobustCell: A Model Attack-Defense Framework for Robust Transcriptomic Data Analysis

**DOI:** 10.1101/2024.11.19.624294

**Authors:** Tianyu Liu, Qi Kang, Yijia Xiao, Xiao Luo, Hongyu Zhao

**Affiliations:** Interdepartmental Program in Computational Biology & Bioinformatics, Yale University, New Haven, 06511, CT, USA; Department of Biostatistics, Yale University, New Haven, 06511, CT, USA; Department of Computer Science, Northeastern University, Boston, 02115, MA, USA; Department of Computer Science, University of California, Los Angeles, Los Angeles, 90095, CA, USA

**Keywords:** Trustworthy Deep Learning, Model Attack, Model Defense, Graph Neural Network, Single-Cell Transcriptomics, Spatial Transcriptomics

## Abstract

Computational methods should be accurate and robust for tasks in biology and medicine, especially when facing different types of attacks, defined as perturbations of benign data that can cause a significant drop in method performance. Therefore, there is a need for robust models that can defend attacks. In this manuscript, we propose a novel framework named RobustCell to analyze attack-defense methods in single-cell and spatial transcriptomic data analysis. In this biological context, we consider three types of attacks as well as two types of defenses in our framework and systemically evaluate the performances of the existing methods on their performance of both clustering and annotating single cells and spatial transcriptomic data. Our evaluations show that successful attacks can impair the performances of various methods, including single-cell Foundation Models. A good defense policy can protect the models from performance drops. Finally, we analyze the contributions of specific genes toward the cell-type annotation task by running the single-gene and group-genes attack methods. Overall, RobustCell is a user-friendly and extension-flexible framework for analyzing the risks and safety of analyzing transcriptomic data under different attacks.

## Introduction

Machine Learning, especially deep learning, is widely used in processing and analyzing biological data [1–3]. However, such methods may be vulnerable to model attack for both white-box models and black-box models [4–7]. Toxic attacks including perturbations and noise on the training data and testing data can significantly degrade the model performance, thus affecting their deployment [8]. To dive deep into the problems caused by toxic attacks, defense methods have been developed to protect the models under different types of attack to preserve their attack-free performances [4, 9]. For biomedical research and clinical applications [10–12], model robustness is important, especially in the analysis of sequencing data [13, 14]. However, regarding the systematic discussion of attack and defense for omics including single-cell RNA sequencing (scRNA-seq) data [15, 16] and spatial transcriptomic data [17, 18], there is limited study. The problem of attacks on omics is important, as (i) They are naturally occurring, such as data noise and batch effect; and (ii) Attacks represent emerging risks as machine learning models become increasingly deployed in clinical and biomedical settings, and attacks on omics are hard to detect as well as defended. In this manuscript, we focus on these two specific types of sequencing data for our analysis of robustness. scRNA-seq are represented by matrices with rows corresponding to cells and columns corresponding to genes, while spatial transcriptomic data with rows corresponding to spots and columns corresponding to genes, plus spatial coordinates for spots. Spots can be cells (such as Xenium-based data [19]) or niches of cells (such as Slide-seq V2-based data [20]). Many machine learning methods have been developed to analyze scRNA-seq and spatial transcriptomic data [21–23], and it is important to study the stability of these methods under different attacks.

Although robustness analysis has achieved great success in handling system, image or text data [4, 24], we cannot directly apply these methods to sequencing data due to the following gaps. The first gap is the domain difference. Methods for attacking or defending widely discussed in computer vision and natural language processing cannot handle the sequencing data [5, 7], and formats with sequencing data are also more noisy [25]. The second gap is the simulation of model attack. Although [26] considered two attack methods at the gene level for cell annotation, their methods did not consider correlations among genes. As well-documented in gene co-expression networks [27, 28] and gene regulatory networks [29, 30], the change of expression of one gene correlated with other genes in its network(s). [31] discussed the application of computational attack methods for disease state classification using single-cell data. However, this work focuses on methods (including ResNet [32] and ViT [33]) in the area of computer vision instead of those used in bioinformatics. Similarly, [34] only discussed the effect of backdoor attack for single-cell Foundation Models, which are not widely deployed. The third gap is the choices of tasks. Cell-type or cell-state annotation as well as cell clustering are common tasks in sequencing data analysis, yet there has not been a systematic study to explore and evaluate robust learning for these tasks. The fourth gap is the development of model defense methods. [35] explored the contribution of adversarial training as a defense method for the attack targeting cell-type annotation, and [36] discussed a closed-resource model for the attack targeting cell clustering. Despite these efforts, model defense has not to be systematically analyzed. The final gap is the robust analysis of spatial transcriptomic data. Graph neural networks (GNNs) are widely used in the study of spatial transcriptomic data [37–39], but the robustness of GNN-based methods for spatial data analysis has not been studied. In summary, the robustness of single-cell analysis tools is an under-explored but important and challenging area, and the goal of this paper is to fill in these gaps.

In this manuscript, we introduce a novel framework for analyzing the attack and defense methods for three well-defined tasks in single-cell and spatial data analysis, including cell-type annotation, cell clustering, and spot-type annotation. To unify the approaches for model attacks, we propose and evaluate the computation-oriented attack and the biology-oriented attack, for these tasks. The computation-oriented attack includes noise-based attacks targeting both expression matrices and graphs, while the biology-oriented attack includes perturbations and the consideration of diseased cells. As a framework for analyzing model robustness, we also include and evaluate current methods for model defense in single-cell data analysis. Overall, we offer a useful tool for researchers interested in the robustness of single-cell data analysis to build a trustworthy machine learning algorithm for the cell-level or spot-level data analysis.

## Results

### Overview of RobustCell

RobustCell has methods for both model attack and model defense. We consider both computation-oriented, biology-oriented, open-box, and closed-box methods to attack single-cell data classifiers. We also analyze the effects of such attacks on cell clustering. Moreover, we consider graph-based methods to attack classifiers for spatial transcriptomics. The poisoned datasets are defined as datasets attacked by these methods, and a dataset that has not been attacked is defined as a clean dataset. Moreover, we consider different defense methods to defend the base models against these three attacks. In addition to analyzing the changes in task-related metrics brought by different attack methods, we also discuss the impact of model attacks on the biological structure of single-cell data, especially for marker genes of different cell types. Our framework is summarized in Figure 1.

**Fig. 1.**
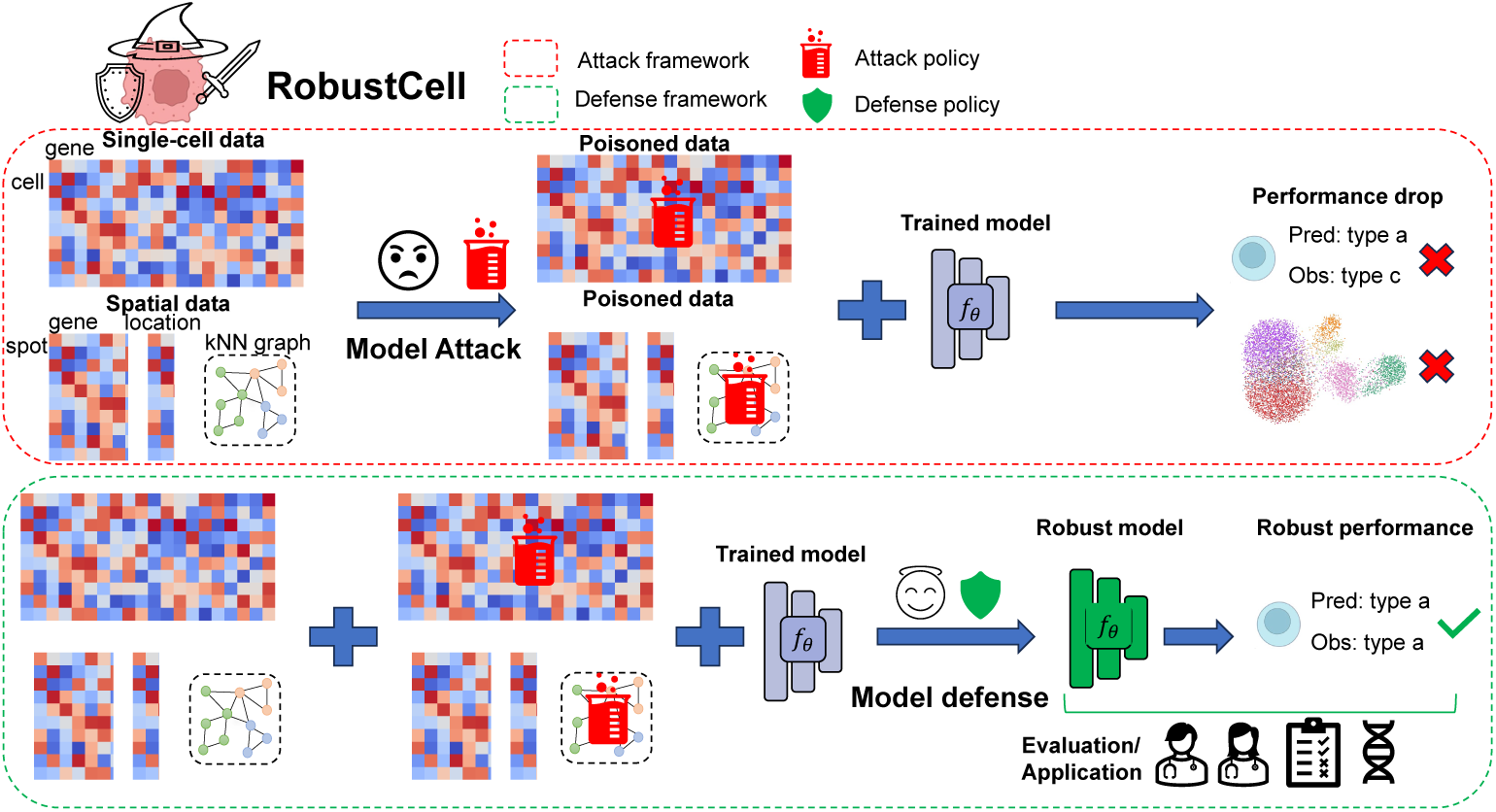
The landscape of RobustCell. The red block represents our framework for constructing and evaluating model attack methods, and the green block represents our framework for constructing and evaluating model defense methods. Here, we consider scRNA-seq datasets (as single-cell data) and spatial transcriptomic datasets with their k-nearest-neighbor (kNN) graphs from locations of spots (as spatial data and kNN graphs). *f_θ_* represents a model for cell-type or spot-type annotation. After having the experimental results for both model attack and model defense, we will investigate the insights from analyzing the model attack-defense framework for applications in single-cell biology.

### Effect of different attack methods on the performance of cell-type classifiers

We first investigated the effect of different attack methods for classifiers focusing on cell-type annotation. By analyzing three different scRNA-seq datasets, denoted as Pancreas [40–43], Aorta [44] and PBMC [45–48], we evaluated both computation-oriented attack methods and biology-oriented attack methods. The computation-oriented attack methods include random attack, Projected Gradient Descent (PGD) attack [49], Fast Gradient Sign Method (FGSM) attack [50], Deepfool attack [51], ensemble attack [52], and composite attack [53, 54]. Random attack means we inject random noise from a Gaussian distribution to the original gene expression profiles. FGSM attack and PGD attack mean we inject gradient-based noise into the original gene expression profiles. Deepfool attack means we iterate adjusting the expression profiles and stop when the minimal change that alters the model output is found. Moreover, we also consider different approaches to aggregate different attacks. Ensemble attack means we average the results from different attacks and composite attack means we adjust the sequence of different attack methods by the greedy-searching approach based on accuracy. The biology-oriented methods include the perturbation attack and the disease attack. Perturbation attack means we use the gene expression profiles from perturbed cells as input for classifiers trained by normal cells, and the perturbation information is arbitrary as we consider all of the measured perturbed gene expression profiles from the selected dataset. Disease attack means we use the gene expression profiles from diseased cells based on their phenotypes as input for classifiers trained by normal cells (known as base classifiers). Our experiments were developed in such a setting: we trained a classifier without attack to generate a base classifier and evaluated the performances of different attacks based on the metrics for classification (Accuracy, (weighted) Precision, (weighted) Recall, and (weighted) F1-score) by using the noisy data and the base classifier. Based on a benchmarking research [55], we included a Support Vector Machine with a rejection setting (*f_θ_* = *SV M_rej_*) as a representative method based on machine learning; moreover, we considered a neural network for classification (*f_θ_* = *Cal*) [35, 56] as a representative method based on deep learning; and scGPT [57] and Geneformer [58] as examples of pre-trained single-cell Foundation Models, since these two models performed well in annotating cell types reported by [59] and pre-training stage might offer more prior information to create a more robust classifier [60]. In our experiments, we follow the default setting of fine-tuning selected models and keep the splitting of cells in each dataset as 60% for training, 10% for validation, and 30% for testing.

We first evaluated the effect of computation-oriented attack methods. In Figure 2 (a), we visualize the distribution of cell types by running Principal Component Analysis (PCA) [61, 62] followed by Uniform Manifold Approximation and Projection (UMAP) [63] for dimension reduction. Figures 2 (b)-(d) represent the comparisons among the default setting and attack methods under a different parameter *eps* based on four different classification metrics. Here, the classifier is *Cal*, and we used the Pancreas dataset. According to these figures, all attack methods can decrease the performance of *Cal*. For the other classifier *SV M_rej_*, we visualize the experimental results in Supplementary Figures 4 (a)-(c), with a similar conclusion from these figures. An ideal attack method should both significantly reduce the model’s performance and be challenging to identify. For example, by increasing the noise level of the random attack, clusters of cells become less easily separated, and the performance of *Cal* becomes worse than the clean mode, according to Figure 2 (b). These figures show that the impact of the attack produced by the PGD method is the most obvious, as the classifier mis-classified all the samples with incorrect cell types. Moreover, there is an upper bound on the effect of the FGSM method, which means that the effect of the FGSM method tends to stay the same when *eps >* 5. In addition, according to Figure 2 (e), as an overall comparison across different datasets, the Deepfool method is also very effective. The Deepfool method is less likely to be detected than PGD because the cell-type distribution in the UMAPs to which the Deepfool method corresponds is easily distinguishable, and thus, its effect on the data is less likely to be detected. In addition, we do not find a clear advantage of ensemble attack or composite attack based on the accuracy shown in this figure, so the single-type attack is more effective for scRNA-seq data.

**Fig. 2.**
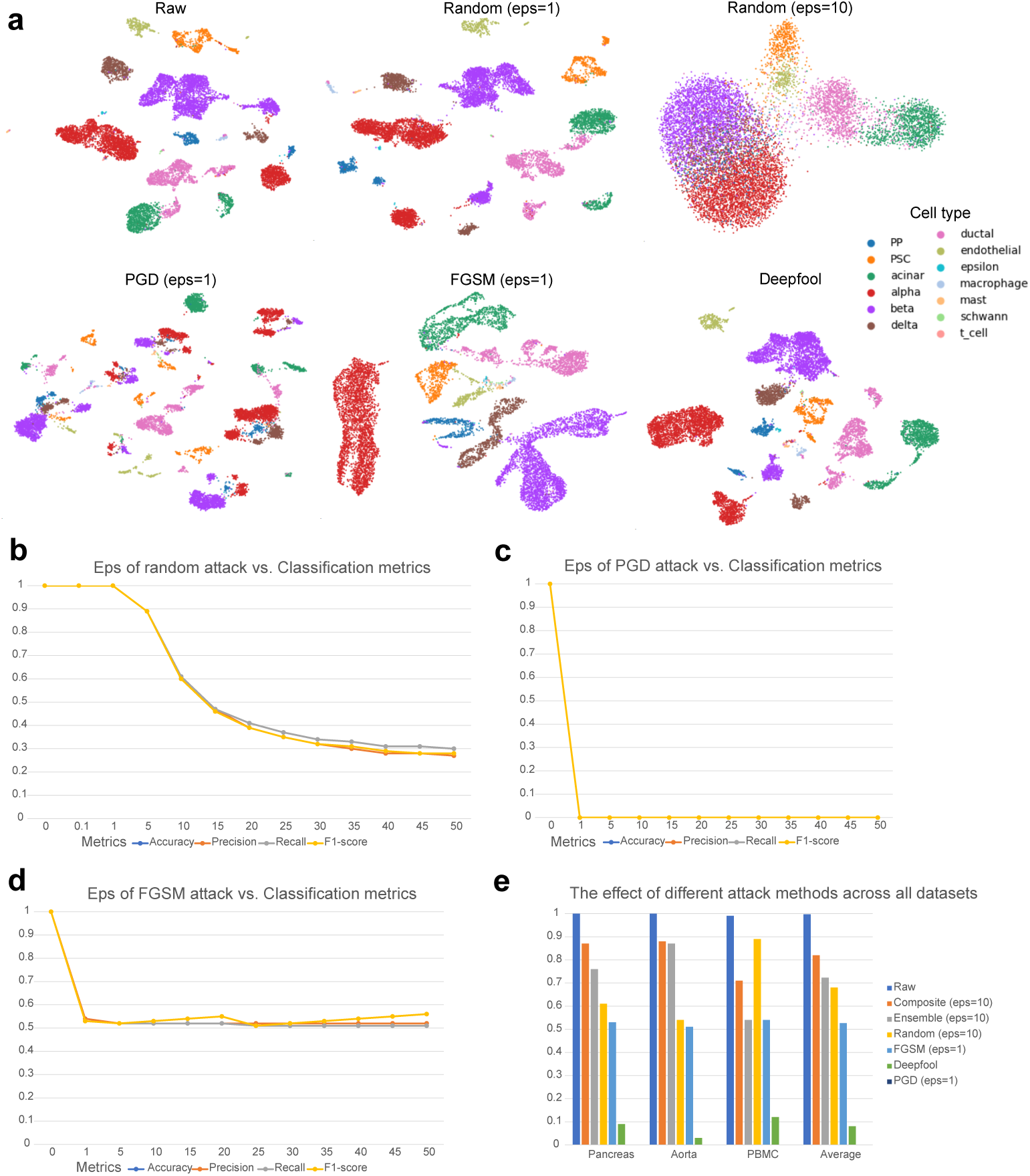
Evaluations for the computation-based attack methods. The term *eps* represents the hyper-parameter used to control the strength of the attack for different methods. Higher *eps* represents a stronger attack. (a) The UMAPs of gene expression profiles from the Pancreas dataset before and after different attack methods. (b) The relationship between the strength of random attack and the scores of different classification metrics for the Pancreas dataset. (c) The relationship between the strength of the PGD attack and the scores of different classification metrics for the Pancreas dataset. The relationship between the strength of the FGSM attack and the scores of different classification metrics for the Pancreas dataset. (e) The effect of different attack methods on cell-type classification performances across different datasets.

In Supplementary Figures 4(a)-(c), we report the performance of scGPT under different attack strategies across the three evaluated datasets, and in Supplementary Figures 4(a)-(c), we present the corresponding results for Geneformer. Overall, we observed that both foundation models were vulnerable to poisoned input data, with adversarial or corrupted perturbations consistently leading to reduced prediction accuracy compared with clean-data settings across all the attack methods. This suggests that, although scGPT and Geneformer are pretrained on large-scale single-cell datasets and can capture rich biological representations, their downstream predictions remain sensitive to distributional shifts and maliciously modified gene-expression profiles. The performance degradation was observed across multiple datasets and attack types, indicating that this vulnerability is not specific to a single model architecture or dataset but reflects a broader robustness challenge for single-cell foundation models. These results further support the need for systematic robustness evaluation when deploying such models in biological analysis pipelines, particularly in settings where input data quality may be uncertain or where corrupted samples could affect downstream biological interpretation. Thus, our findings highlight that poisoning attacks can compromise not only task-specific models but also large pretrained single-cell foundation models, emphasizing the importance of developing effective detection, defense, and reliability-assessment strategies for secure single-cell foundation models.

We then evaluated the effect of biology-oriented attack methods. Figures 3 (a)-(b) display the distribution of conditions for the Aorta dataset and the Openproblems dataset [64]. The former dataset is an example used for performing the disease attack, while the latter is used for performing the perturbation attack. We trained classifiers based on datasets under the control condition and evaluated the classifiers using cells with non-control conditions. According to Figures 3 (c) and (d), the disease attack method under different diseases successfully affects the robustness of both *Cal* and *SV M_rej_*. Moreover, *SV M_rej_* is more robust than *Cal* under this attack since its performance drop was less noticeable. Based on Figure 3 (e), the perturbation attack method also successfully weakened the capacity of *Cal*. Such observation is consistent for *SV M_rej_*, shown in Supplementary Figure 4. In addition, the biology-oriented attack methods also lead to the performance drop of scGPT and Geneformer, shown in Supplementary Figures 4 (d)-(e) and Supplementary Figures 4 (d)-(e). Pre-training might not be enough to have a robust single-cell classifier.

**Fig. 3.**
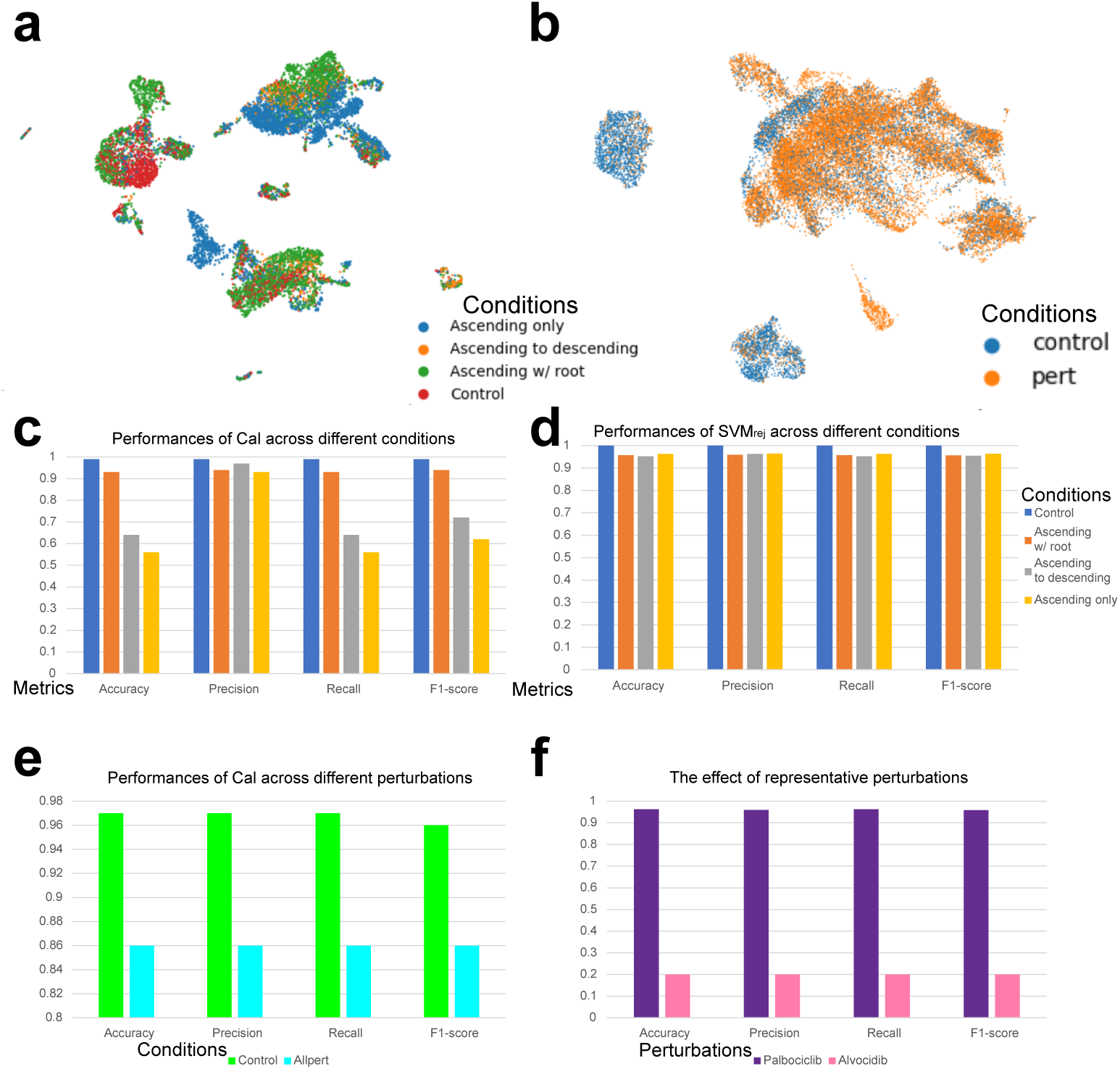
Evaluations for the biology-based attack methods. (a) The UMAPs of gene expression profiles from the Aorta dataset colored by different conditions. (b) The UMAPs of gene expression profiles from the Openproblems dataset colored by different conditions. (c) The effect of the disease attack method on cell-type classification performances of *Cal* based on the Aorta dataset. (d) The effect of the disease attack method towards cell-type classification performances of *SV M_rej_* across different datasets. (e) The effect of the perturbation attack method towards cell-type classification performances of *Cal* based on the Openproblems dataset. (f) The effect of representative perturbations towards cell-type classification performances of *Cal* based on the Openproblems dataset.

Since the Openproblems dataset contains 141 drug-based perturbations, it would be interesting to investigate whether there is a difference in the effect of different perturbations on the classifiers. Therefore, we record the results of classification metrics across all the perturbations in Supplementary Data 1. We also select the perturbations with the highest and lowest accuracy, shown in Figure 3 (f). From this figure, we could infer that Alvocidib might strongly affect the biological systems in different cells, while Palbociclib might not affect different cells much. Since we ensured cell types after the attack could still be found in the control case, our results were not confounded by the possible distribution shift in the cell-type level. Therefore, our study demonstrated that biology-oriented attack methods can also explain differences in biological systems across cell conditions.

### The performance of cell clustering can be used to analyze a successful attack’s identifiability

We expected that a good attack method should lead to obvious performance drop in classification but not in clustering. Therefore, to analyze the effect of model attack methods on cell clustering and the identifiability of these attack methods, we also investigated the performances of cell clustering after attacking these three datasets used in the previous section. Such results are summarized in Figure 4. The metrics we used for evaluating cell clustering included Normalized Mutual Information (NMI), Adjusted Rand Index (ARI), and Average Silhouette Width (ASW) referred from [65]. We also averaged their values for an overall comparison. We first analyzed the effect of *eps* On different attack methods for cell clustering. Increasing *eps* means that we expect to have a stronger attack. According to Figures 4 (a)-(c), a stronger random attack or a stronger PGD attack both led to a performance drop in cell clustering. A strong positive correlation exists between the effect of the random attack method for cell-type annotation and for cell clustering (Pearson Correlation Coefficient=0.91, p-value=1.1e-5, for Accuracy vs. Averaged clustering metrics of the Pancreas dataset). The PGD attack is easier to identify than the random attack because it leads to a sharper performance drop in cell clustering. However, the strength of the FGSM attack did not affect the clustering performance much, and it even led to slight improvement after such an attack on the Pancreas dataset. We have similar conclusions for the two datasets, shown in Supplementary Figures 4 (a)-(f). Considering the error bound of this attack for the cell-type annotation task, we believe this approach is not a very successful attack method. We also included Deepfool and other attack methods under their representative settings in our discussion, shown in Figures 4 (d)-(f). From these figures, the Deepfool method does not affect the performances of cell clustering too much. However, it can significantly affect the performance of classifiers for cell-type annotation. The effect of Deepfool attack cannot be directly detected by clustering cells, and it still confuses cell-type classifiers. Overall, the analysis of the cell clustering results after the attack helps us to understand the characteristics of the different attacks and the extent of their impact on the negative effects of selected models.

**Fig. 4.**
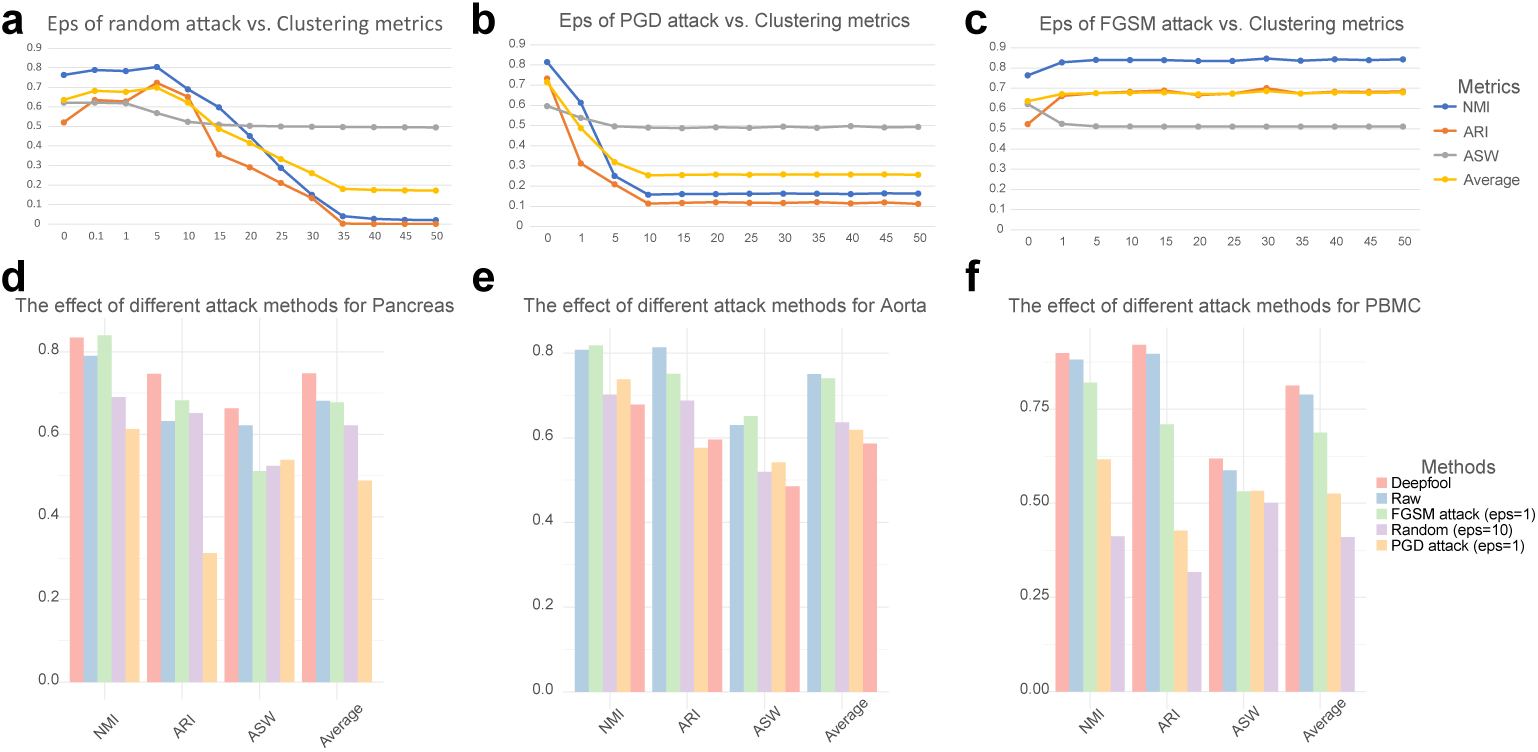
Investigating the performances of cell clustering under different attack methods. (a) The relationship between the strength of random attack and the scores of different clustering metrics for the Pancreas dataset. (b) The relationship between the strength of the PGD attack and the scores of different clustering metrics for the Pancreas dataset. (c) The relationship between the strength of the FGSM attack and the scores of different clustering metrics for the Pancreas dataset. (d) The effect of different attack methods towards cell clustering of the Pancreas dataset. (e) The effect of different attack methods towards cell clustering of the Aorta dataset. (f) The effect of different attack methods towards cell clustering of the PBMC dataset.

### Effect of different attack methods on the effectiveness of graph-based spot-type classifiers

In this section, we analyzed graph-based model attack for spot-level data analysis [6]. The target of such an attack is the edge of graphs. By perturbing the edges in a graph dataset, it is possible to reduce the performance of GNNs on the node classification task [66]. Different from single-cell datasets, spatial datasets contain extra spatial information, and thus, such information can be used to help identify the types of spots [37]. We can treat spots as nodes and construct neighbor graphs based on the distances computed from the locations of spots. The number of neighbors ranges from 10 to 50, and thus we can capture different levels of node modeling. The larger neighbor number is, the more edges we can create. To support the advantages of using GNNs, we utilize the Starmap dataset as an example and visualize the distribution of cell types by locations in Supplementary Figure 4 (a). By comparing Graph Convolutional Networks (GCNs) [67] with Multi-layer Perceptrons (MLPs) shown in Supplementary Figure 4 (b), we demonstrate that using GCNs can more accurately predict spot types. To evaluate the strength of different attack methods and the robustness of GNN-based methods for the spot-type classification task, we evaluated the performances of four different attack methods (including meta attack [68], DICE attack [69], graph random attack [66], and Minmax attack [70]) by using GCNs under two different spatial transcriptomic datasets. The metrics used in this section are the same as those for evaluating single-cell classifiers. Details of different attack methods are summarized in the Methods section.

In Figures 5 (a) and (b), we display the distribution of spot types for the Slide-seq v2 dataset (left) [20] and the Xenium dataset (right) [71]. From these figures, we found that some spot types have spatially variable patterns, for example, C1 C2 C3 Subiculum from the Slide-seq v2 dataset and ECM1+ Malignant from the Xenium dataset. This observation suggests that disruption of neighborhood relations may affect the accuracy of the spot-type classifier. Considering the influence of *k* for constructing the kNN graphs, in Figure 5 (c), we visualize the relationship between the value of *k* and the classification performance of our GCN classifier based on the Slide-seq v2 dataset. According to this figure, increasing the number of neighbors used to construct the graph leads to performance drop of our GCN classifier. This observation can be partially explained by class homophily score [72, 73] as a type of graph homophily score [74]. Class homophily score measures the contribution of edges in a graph to the node classification task. Increasing the number of edges can introduce more information in the training process, while the homophily score will decrease, and thus may affect the model performances. Our conclusions are further supported by Supplementary Figures 4 (b) and (c). All datasets show a negative correlation between the number of neighbors and classification performances with GCN, and the Starmap dataset shows an upward and then downward trend for spot-type classification. Furthermore, we show the results of different graph-based attack methods by averaging the classification metrics across different *k* based on the Slide-seq v2 dataset in Figure 5 (d). The strongest attack method for this dataset is meta attack. The contributions of the DICE attack and the graph random attack are not obvious compared with other methods. Interestingly, perturbing the kNN graph based on the Minmax attack method leads to better classification performance shown in Figure 5 (d). We, therefore, argue that algorithms that attack graphs produce more complicated results than algorithms that attack gene expression profiles for the classification task. For the Xenium dataset, we display figures with the same settings in Figure 5 (e) for the relationship between *k* and classification performance and in Figure 5 (f) for the effect of different attack methods. Based on these two figures, we have similar conclusions to the Slide-seq v2 dataset results. Overall, introducing noise into the kNN graph might help predict spot types. However, the Meta attack method consistently decreases the performance of our GCN classifier under different datasets. Based on our evaluations, it is the most suitable attack method for kNN graphs.

**Fig. 5.**
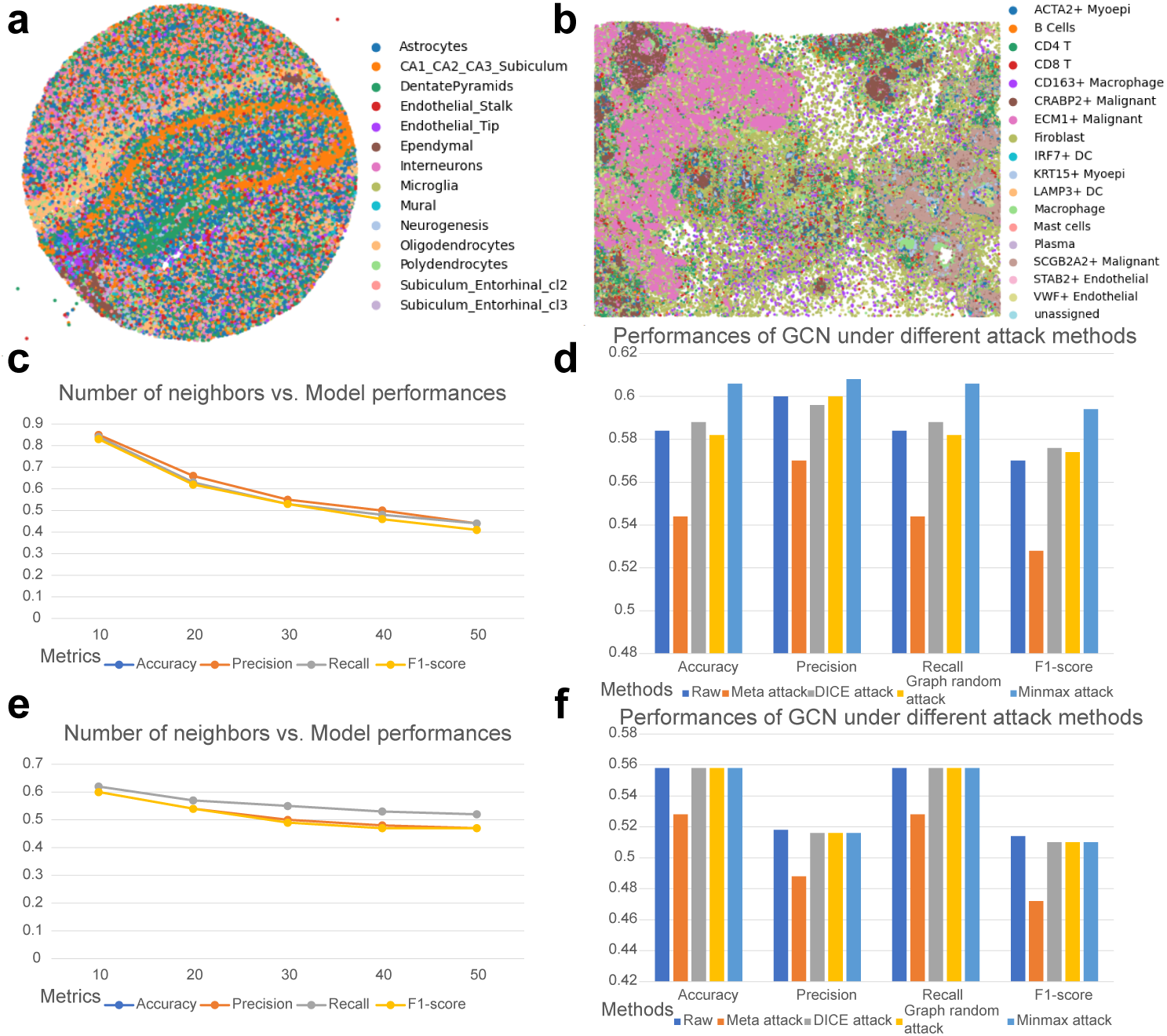
Evaluations for the graph-based attack methods using spatial transcriptomic datasets. (a) The plot of the cell-type distribution of the Slide-seq v2 dataset. (b) The plot of cell-type distribution of the Xenium dataset. (c) The plot of the relationship between *k* and classification performances is based on the Slide-seq v2 dataset. (d) The plot of the classification performances based on different attack methods using the Slide-seq v2 dataset. The results are averaged by different *k* or each method. (e) The plot of the relationship between *k* and classification performances based on the Xenium dataset. (f) The plot of the classification performances based on different attack methods using the Xenium dataset. The results are averaged by different *k* for each method.

### Evaluations of model defense methods

After evaluating different attack methods for single-cell and spot-level classifiers, we investigated the possibility of discovering a set of suitable policies for model defense. Our goal is to find suitable defenses against different attack methods, thus reducing the attack algorithms’ negative impact on the classifiers’ effectiveness. For the computation-oriented attack methods and biology-oriented attack methods, we considered three approaches for model defense, including marker gene selection [75], adversarial training [35], and composite defense. The idea of marker gene selection is to train a classifier based on a clean data subset by marker genes. We used the Wilcoxon rank-sum test based on Scanpy [76], and 40 marker genes per cell type suggested by [46]. The idea of adversarial training is to train a classifier based on clean and poisoned data. We also considered the composite version of these two approaches, composite defense. For the graph-based attack methods, we considered four approaches for model defense, including GCNSVD [77], GCN-Jaccard [78], RGCN [79], and SimPGCN [80]. Graph defense methods aim to train a GNN with robust structures or policies based on poisoned graphs. Details of all the model defense methods are summarized in the Methods section, and all the evaluations here are based on the poisoned data unless otherwise specified.

For the defense of computation-oriented attack methods, we visualize the performances of different defense methods in Figures 6 (a) and (b). Figure 6 (a) shows the comparisons between the metrics of attack methods and the metrics of the corresponding marker gene selection method for the cell-type annotation task. Based on this figure, marker gene selection as a defense method successfully increased the performances of our classifiers under attack methods, including Deepfool, PGD, and FGSM, across all kinds of datasets. We also tested the relationship between the number of highly-variable genes (HVGs) and the performances of different attack methods to preclude gene number as a confounder, shown in Supplementary Figure 4 for the Pancreas dataset, where the figure shows that reducing the number of HVGs cannot consistently defend the attacks. Therefore, only the marker gene selection can work as a defense method. However, marker gene selection cannot be used to defend poisoned data from the random attack method. Figure 6 (b) shows the comparisons between the metrics of attack methods and the metrics of the corresponding adversarial training method. From this figure, the adversarial training method generated better performance on cell-type annotations under all types of attack methods across all datasets. Regarding the composite defense, we show the results in Supplementary Figure 4, and the composite defense method performs better than adversarial training in defending the PGD attack and the random attack. Therefore, the performance of the composite defense method is driven by choices of attacks. We also tested the contributions of these two defense methods for *SV M_rej_*, shown in Supplementary Figure 4 for defending the random attack. We found that the adversarial training method can also benefit the robustness of *SV M_rej_*. Therefore, we believe that adversarial training is a crucial method to defend models for robust cell-type classification.

**Fig. 6.**
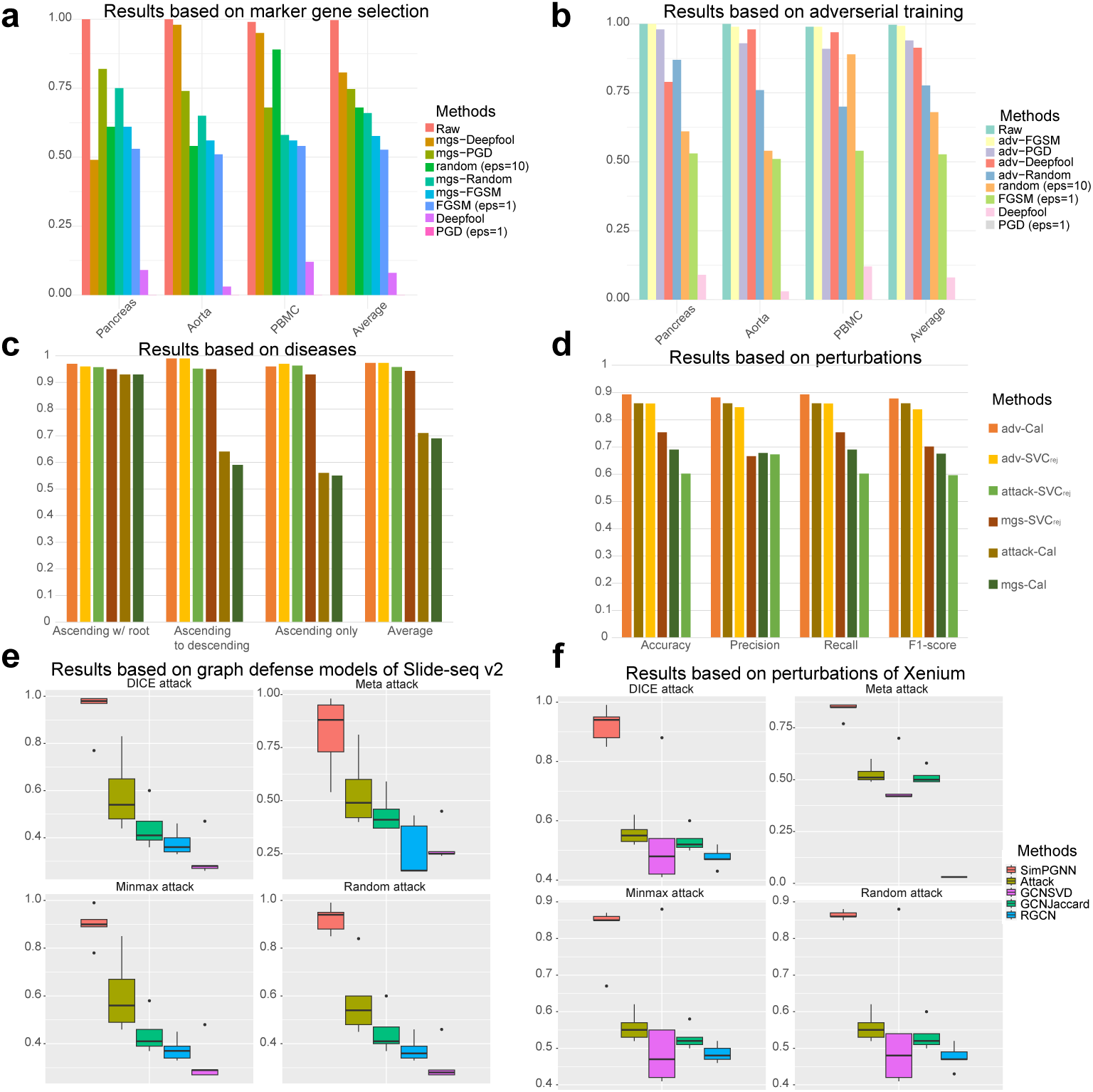
Evaluations of defense methods for different types of attack methods. Here *Raw* represents the accuracy under the clean dataset. Defense methods starting with *mgs* represent that these attack methods are defended by the marker gene selection method, and defense methods starting with *adv* represent that these attack methods are defended by the adversarial training method. (a) The comparison of the accuracy of original attack methods and the corresponding defense results using marker gene selection across different datasets. (b) The comparison of the accuracy of original attack methods and the corresponding defense results using adversarial training across different datasets. (c) The comparison of accuracy between original attack methods and the corresponding defense results across different types of diseases. (d) The comparison of four different metrics between original attack methods and the corresponding defense results under the perturbations. (e) The comparison of accuracy between original graph-based attack methods and the corresponding defense results based on the Slide-seq v2 dataset. (f) The comparison of accuracy between original graph-based attack methods and the corresponding defense results based on the Xenium dataset.

We also investigate the reliability and conditions under which marker gene selection works. Considering the dataset from human pancreas, we have added three attack methods: 1. Normal noise added for genes except marker genes (non-marker noise); 2. Normal noise added for marker genes (marker noise); 3. Gene expression dropout attack for maker genes (marker dropout). The classifier is SVC. The results are updated in Table 1. We can interpret the results in two ways. First, when the attack occurs in non-marker genes, the marker-gene classifier remains unaffected, indicating that this method can indeed serve as a defense against feature selection. Second, when attack occurs in the marker genes themselves, the marker-gene-based classifier rapidly breaks down, supported by an obvious performance drop. Therefore, this defense method can only support the condition where marker genes are not directly affected. We are also aware of the limitations of this method, so in the future, we may explore ways to expand upon it, such as using distribution similarity to assign a score to the likelihood of a gene being a marker; if a marker gene falls below a certain score, we would not recommend using this method.

**Table 1.** Marker Gene Selection Experiments Under Various Conditions.

| Feature Set | Attack Type | $\epsilon$ | Accuracy | Macro-F1 |
| --- | --- | --- | --- | --- |
| All genes | Clean | 0 | 0.974 | 0.846 |
| Marker genes | Clean | 0 | 0.964 | 0.722 |
| All genes | Non-marker noise | 10 | 0.714 | 0.491 |
| Marker genes | Non-marker noise | 10 | 0.964 | 0.722 |
| All genes | Marker noise | 10 | 0.024 | 0.053 |
| Marker genes | Marker noise | 10 | 0.007 | 0.023 |
| All genes | Marker dropout | 1 | 0.430 | 0.334 |
| Marker genes | Marker dropout | 1 | 0.239 | 0.039 |

For the defense of biology-oriented attack methods, we visualize the performance of different defense methods in Figures 6 (c) and (d). Regarding the contributions of these two defense methods for defending the disease attack method, we show the experimental results in Figure 6 (c), and found that adversarial training improved the performances of both *SV M_rej_* and *Cal* for classification. However, marker gene selection did not contribute to defending against this type of attack. Regarding the contributions of these two defense methods for defending the perturbation attack method, we display the experimental results in Figure 6 (d). We found that adversarial training improved the performances of both *SV M_rej_* and *Cal* under this type of attack. In Supplementary Figures 4 (a)-(b), we illustrate the comparisons between the composite defense method and adversarial training method for the Aorta dataset and the Openproblems dataset, and we do not find the superiority of the composite attack. Therefore, the adversarial training method is still a better choice to defend against attacks from disease or perturbations.

For the defense of graph-based attack methods, we display the performances of different defense methods in Figures 6 (e) and (f) by different datasets. In Figure 6 (e), we found that SimPGCN consistently performed well across different attack methods for the Slide-seq v2 dataset. In contrast, other defense methods could not improve the classification performances for spot-type classification. We also had similar conclusions based on our investigation for the Xenium dataset, shown in Figure 6 (f). Therefore, SimPGCN is the only method we recommend for defending graph-based attacks.

Overall, our study shows that adversarial training can effectively defend against various attacks on cell-type classifiers with multi-layer perceptrons (MLPs) as base models, while SimPGCN can effectively defend against various attacks on spot-type classifiers with GCNs as base models.

### An investigation of whether model attacks affect the marker genes of single-cell datasets

Since the marker gene selection approach can be used to defend certain attack methods for single-cell classifiers, we are interested in discussing the effect of model attacks towards the clean single-cell gene expression profiles, as well as the contribution of each marker gene towards the classification performance. We used the Wilcoxon rank-sum test to discover marker genes of each cell type, and the number of marker genes for each cell type is still 40.

In Figures 7 (a) and (b), we visualize the comparisons of marker genes discovered based on the clean gene expression profiles and the gene expression profiles poisoned by the computation-oriented attack methods. The single-cell dataset here is the Pancreas dataset. In Figure 7 (a), we show the overlap of marker genes discovered by the Wilcoxon rank-sum test under different cases. The profiles processed by the PGD method have the largest marker gene overlap ratio compared with the marker gene list from the clean dataset. In contrast, the profiles processed by the Deepfool method have the smallest overlap ratio. As a result, Deepfool-based attacks can largely change the classifier’s perception of different cell types. Furthermore, we investigated the change of the Gene Ontology (GO) enrichment analysis [81, 82] results before and after attacking for the same dataset, shown in Figure 7 (b). The results were computed based on the ratio between the number of significant pathways and the number of all pathways involved in the marker gene list. After ranking the results by the averaged values across cell types, we found that Deepfool-based methods also maximally altered the extent of enrichment of marker genes in different cells. However, after performing the Wilcoxon test between the enrichment ratios from clean profiles and the enrichment ratios from the poisoned profiles, we did not identify significant attack methods for affecting the GO enrichment level (all p-values are greater than 0.05). Therefore, cell-type heterogeneity is a factor that must be considered, and the effect of attack methods for different cell types is not in the same direction. Moreover, we notice that after certain attacks, T cells, alpha cells, epsilon cells, and schwann cells show zero enrichment ratios, which implies that these cell types are easier to be affected by computation-oriented attacks.

**Fig. 7.**
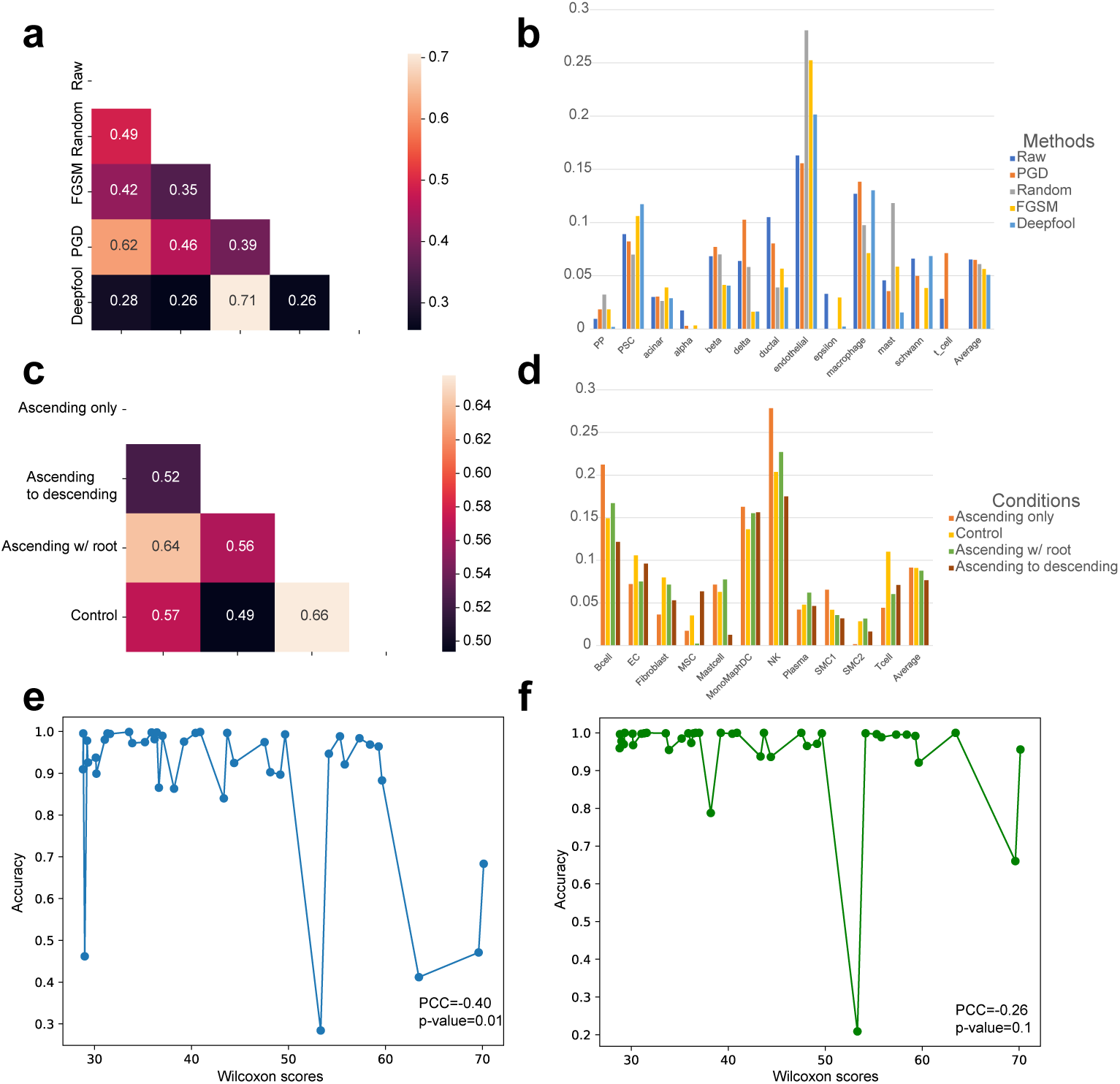
Results of the relationship between marker genes and model robustness. (a) The heatmap colored by the overlap of marker genes among the clean gene expression profile and the poisoned gene expression profiles for the Pancreas dataset. The *eps* settings are the same as our previous representative choices. (b) The ratio of GO enrichment analysis among different cell types discovered based on the clean gene expression profile and the poisoned gene expression profiles based on the Pancreas dataset. (c) The heatmap colored by the overlap of marker genes among the clean gene expression profile and the poisoned gene expression profiles for the Aorta dataset. (d) The ratio of GO enrichment analysis among different cell types discovered based on the clean gene expression profile and the poisoned gene expression profiles based on the Aorta dataset. (e) The relationship between the Wilcoxon score of the gene targeted by the pos attack and the accuracy of the corresponding classifier. The genes are ranked by their Wilcoxon scores in ascending. (f) The relationship between the Wilcoxon score of the gene targeted by the neg attack and the accuracy of the corresponding classifier. The genes are ranked by their Wilcoxon scores in ascending.

In Figures 7 (c) and (d), we visualize the comparisons of marker genes discovered based on the clean gene expression profiles and the gene expression profiles poisoned by the biology-oriented attack methods. The dataset we used here is the Aorta dataset, and the attack method is a disease attack. We show the overlap ratio of marker genes before and after attacking in Figure 7 (c), and the profiles under the condition *Ascending w/ root* have the largest overlap ratio comparing with the profiles from the condition *Control*, while the profiles under the condition *Ascending to descending* have the lowest overlap ratio. Moreover, by performing the Wilcoxon rank-sum test for the GO enrichment ratios shown in Figure 7 (d) between the ratios from the condition *Control* and the ratios from other diseased conditions, *Ascending to descending* is the only condition which has significant level lower than 0.1 (p-value=0.083). All p-values can be found in Supplementary Data 2. Therefore, different diseases also tend to affect the marker genes across cell types differently.

Finally, to investigate the contributions of different marker genes for the cell-type classifier *Cal*, we implemented the single gene attack proposed by [35] as well as a novel pathway attack to adjust the expression levels of specific genes across all selected cells. For single gene attack, we subset alpha cells from the Pancreas dataset and adjusted each marker gene’s expression levels by multiplying +100 (known as a pos attack) or −100 (known as a neg attack). The scales are suggested by [83]. We visualize the relationship between the testing scores and the accuracy after the pos attack for each gene in Figure 7 (e). The pos attack for marker genes with higher scores also tended to lead to a more significant drop in accuracy, supported by the strong negative Pearson correlation between these two scores (p-value=0.01*<*0.05). However, if we consider the same results but under the neg attack shown in Figure 7 (f), there was no significant Pearson correlation between the same group of scores (p-value=0.1). Therefore, the pos attack can also be used to analyze the contribution of each gene to the performance of cell-type classification and to evaluate the performances of defense methods. Furthermore, marker genes play an essential role in the automatic cell-type annotation methods, and protecting the expression levels of marker genes can also help defend the pos attack. Conversely, if we applied the pos attack and neg attack for genes in the same GO pathway, the neg attack had a more significant effect, shown in Supplementary Figures 4 (a) and (b). Therefore, the neg attack can be utilized to analyze group effects of co-functional genes.

## Discussion

Robustness is an essential factor to be considered when developing models for biomedical applications. It is also crucial to define a unified model attack-defense framework to compare the effects of different attack/defense policies, especially for single-cell data analysis. In this manuscript, we first highlighted the necessity of having robust cell-type and spot-type classifiers and review the contributions and drawbacks in the current research. We then proposed a general framework to analyze and develop attack and defense methods for the classification problems of both single-cell and single-spot data. Our attack methods include computation-oriented attack methods, biology-oriented attack methods, and graph-based attack methods, focusing on different aspects of problems and types of datasets. Moreover, we also discussed the definition of a good attack method by arguing the identifiability of such a method based on the performance of cell clustering using poisoned datasets.

Furthermore, we implemented and evaluated different defense methods for these three types of attack methods and found that adversarial training policy as a promising approach in defending the computation-oriented attack methods and biology-oriented attack methods for single-cell data, while SimPGCN performed the best in preserving the graph-based attacks for spatial data. Moreover, our framework is easy to extend; thus, it is possible to include more advanced defense methods and novel attack methods.

Considering the information we can gain from analyzing the model attack-defense framework for biology discoveries, we examined the effect of attacking the expression levels of marker genes for different cell types towards classification performances. We also found that the effects of attack methods on marker genes of different cell types in different directions. Changes in more significant marker genes led to more significant decreases in model performances after the pos attack and changes in more significant pathway-related genes led to more significant decreases in model performances after the neg attack. Therefore, our study provides new ideas for analyzing important genes of different cell types as well as functions and shows that protecting marker genes is an approach that can be considered when developing defense approaches for cell-type annotation.

Despite the apparent benefits of our framework, it has several challenges and limitations. First, adversarial training requires including attack data in the training dataset and thus does not provide a defense against unknown attacks. The ideal solution is to have a unified defense framework [84]. Second, extra training requires longer time and larger memory usage for classification, which is hard to avoid. Third, a more complicated model architecture might respond differently to the model attack methods, which should be investigated in future research. Fourth, different methods to discover marker genes also have their bias, so it is important to develop an unbiased and robust method for marker gene discoveries. Finally, the attack methods will be more complicated for affecting multi-modal datasets [85]. Therefore, we plan to extend this framework for analyzing multi-modal attack and defense methods [86, 87] in the future.

## Methods

### Problem statement

In this manuscript, we have a single-cell RNA sequencing dataset as 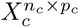, where *n_c_* represents the number of cells and *p_c_* represents the number of genes. Moreover, we have a spatial transcriptomic dataset with 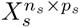 as an expression profile and 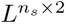 as a location profile. Here *n_s_* represents the number of spots and *p_s_* represents the number of genes. We also construct a kNN graph *G* based on *L*. To annotate cell types or spot types, we have a trained classifier for cell-type classification as *M_c_*(*X_c_*) and a trained classifier for spot-type classification as *M_s_*(*X_s_, G*).

Now consider an attack for the single-cell set as *Attack*(), which transforms the clean datasets into poisoned datasets. The attack towards single-cell datasets can be represented as *X^′^* = *Attack*(*X_c_*) and the attack towards spatial datasets can be represented as *G^′^* = *Attack*(*X_s_, G*). We evaluate the performances of the classifiers under this attack by computing *M_c_*(*X^′^*) and *M_s_*(*X^′^, G^′^*), and then compare the outputs of these models with the observed cell types or spot types.

Regarding the framework of model defense, we consider a model defense method as *Defense*(). We then apply the defense methods for the classifier trained based on datasets specified by the selected method as input. Therefore, we have defense methods as *Defense*(*M_c_*(*X_c_, X^′^*)) for single-cell datasets and *Defense*(*M_s_*(*X_s_, G^′^*)) for spatial datasets. We then compare the outputs of these models with the observed cell types or spot types for evaluation.

By comparing different levels of performance drop under different types of attack methods, we can select the strongest attack method. Moreover, by comparing the different levels of performance preservation under different defense methods, we can select the best defense method.

### Threat model

RobustCell considers adversarial threats against transcriptomic analysis models for cell-type annotation, cell clustering, and spot-type annotation. The adversary’s goal is to degrade model performance by causing misclassification or evasion at inference time, rather than to implant a persistent backdoor or corrupt the full training process.

For single-cell data, the adversary perturbs the gene expression matrix *X_c_* to produce a poisoned matrix *X^′^*. For spatial transcriptomics, the adversary perturbs the spatial kNN graph *G* to produce a poisoned graph *G^′^*. We consider both open-box and closed-box settings. In the open-box setting, the adversary has access to the model architecture, loss function, and gradients, which enables gradient-based attacks such as FGSM, PGD, Deepfool, meta attack, and Minmax attack. In the closed-box or limited-access setting, the adversary does not require model weights or gradients and instead perturbs input data or graph structure through random noise, disease/perturbation shifts, DICE, or graph random attacks. The adversary is not assumed to modify ground-truth labels. Our defense experiments evaluate whether marker-gene selection, adversarial training, composite defense, and graph defense methods can preserve performance under these poisoned inputs.

### Details of different classifiers

For the classifiers trained based on the single-cell datasets, we consider a machine-learning-based classifier *SV M_rej_* and a deep-learning-based classifier *Cal*. *SV M_rej_*represents a support vector machine with rejection option [88, 89]. A support vector machine determines the classification results based on the hyperplanes from all the observations, while the rejection option allows the classifier to not include all the observations. The setup is determined by a Bayes decision rule known as Chow’s rule [90]. Our implementation refers to scikit-learn [62]. *Cal* is a three-layer neural network for the classification task. We utilize the cross-entropy loss [56] and PyTorch lightning framework [91] to train the model.

For the classifier trained based on spatial datasets, we consider a GCN-based classifier *GCN*, including both the message-passing stage and the feature-updating stage. The input data come from the gene expression profiles *X_s_* and a kNN graph *G* generated from *L_s_*. We utilize the cross-entropy loss and the Deeprobust framework to train the model.

### Details of different attack methods

Here we consider three types of attack methods: computation-oriented attack methods, biology-oriented attack methods, and graph-based attack methods. These methods are ranked by the order of appearance in the main text. Computation-oriented attacks are used as stress-test perturbations rather than as fully realistic biological perturbation models. Because directly perturbing the continuous gene-expression matrix may produce biologically implausible profiles, including negative values or disrupted co-expression patterns, these attacks should be interpreted as diagnostic probes of model sensitivity and worst-case vulnerability. In contrast, disease- and perturbation-based attacks provide a more biology-grounded evaluation of robustness under realistic transcriptional shifts. Future work should incorporate biologically constrained adversarial generation, such as non-negativity constraints, preservation of sparsity structure, gene–gene co-expression constraints, pathway-level perturbation priors, and generative models trained on valid single-cell profiles.

Computation-oriented attack methods are inspired by the general attack methods used for image processing. The idea is to inject noise into a clean dataset to create a poisoned dataset. Here *ϵ* is the same as *eps* we used in the main text, and ∇ represents the gradient.

#### Random

The random attack method means we inject the random noise from A Gaussian distribution with 0 as mean and 1 as variance, denoted as *N* (**0**, **1**), to the original gene expression profiles. Here **0** represents a matrix with the same shape as *X_c_* but is filled with all zeros, and **1** represents a matrix with the same shape as *X_c_*but is filled with all ones. This method can be represented as:

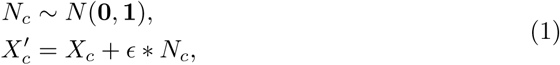

where *ɛ* represents the factor used to control the strength of the random attack and *X^′^* represents the poisoned matrix.

#### Projected Gradient Descent (PGD) [49]

The PGD attack method is based on iteratively changing the entries of the gene expression matrix based on saddle point formulation. Such a method is used to derive the strongest perturbation by optimizing the following equation:

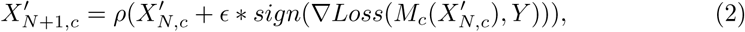

where *ɛ* represents the factor to control the strength of this attack, *ρ*() represents the value of the PGD attack for the matrix *X_c_* with labels *Y*, and *Loss*() represents the loss function. The iteration number is defined as *N* and the initial matrix for the iteration perturbations is *X_c_*.

#### Fast gradient sign method (FGSM) [50]

The FGSM attack is inspired by the backpropagation of optimizing a neural network, and to perform the attack, we can modify the samples with noise in the same direction as the gradient to reduce the model performances. The FGSM attack can be represented as:

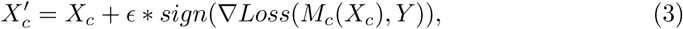

where *ɛ* represents the factor to control the strength of this attack, *Loss*() represents the loss function, and *Y* represents the corresponding labels of the sample matrix *X_c_*.

#### Deepfool [51]

Deepfool is also an iterative attack and it stops when the minimal perturbation which can affect the change of model predicted labels is discovered. For iteration *i*, the decision boundary of a classifier can be represented as a convex polyhedron *P_i_*, that is:

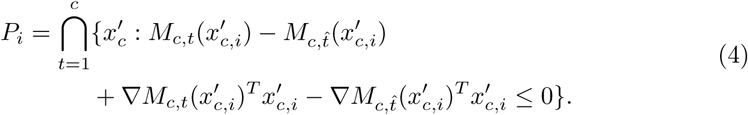

In this definition, *c* represents the number of classes and *t*^^^ represents the observed label before the model attack for the sample *x_c_*. *M_c,t_* represents the model whose output label is *t* given the input example. We start the iteration with *x_c_* and stop the iteration once the perturbed sample *x^′^* reaches the decision boundary. For a single-cell matrix, we combine the perturbed results of each cell to generate the final poisoned matrix *X^′^*. There is no hyper-parameter we need to adjust for this method.

We test the random attack method for both *SV M_rej_* and *Cal* in our experiments, while we only examine the PGD attack, the FGSM attack, and the Deepfool attack for *Cal* because all of these attack methods require the computation of gradients.

Biology-oriented attack methods are inspired by the observations from biology, including the effect of diseases and perturbation towards healthy or control cells. We can access cells with the same proportion of cell types from the tissues under the biology-oriented attack.

#### Ensemble attack [52]

Ensemble attack represents that we average the outputs of different computation-oriented attacks as final outputs after attacking.

#### Composite attack [53, 54]

Composite attack represents that we arrange these four computation-oriented attack methods (including random attack, FGSM attack, PGD attack, and Deepfool attack) in a sequence determined by either the user’s input or a greedy-searching method. The target of the greedy-searching design is to determine the sequence that can generate the lowest classification accuracy for single-cell classifiers in each step.

#### Disease attack

The disease attack represents that we treat the cells under the diseased conditions as input to test the performances of a classifier trained based on healthy cells from the same tissue.

#### Perturbation attack

The perturbation attack represents that we treat the cells under different perturbations as input to test the performances of a classifier trained based on cells under the control case and from the same tissue. That is, the arbitrary perturbation Pert and corresponding expression profiles GEX_Pert_ will be used to replace the normal gene expression profile, and the cell pair will be randomly selected and labels will not change. In this manuscript, we only consider drug-based perturbations. However, we note that our framework also fits with other types of perturbations including datasets from perturb-seq [92].

We test the biology-oriented attack methods for both *SV M_rej_* and *Cal* in our experiments.

The graph-based attack is specifically designed for classifiers trained based on spatial transcriptomic datasets with graph information, and the target of graph-based attack methods is the kNN graph *G* generated from *L*.

#### Meta attack [68]

The meta attack allows users to treat a graph structure matrix as a hyper-parameter and it computes the meta gradient of the given loss function with respect to the original graph structure. Furthermore, the meta attack utilizes a greedy approach to assign perturbations based on the value of the meta gradient. That is, given the meta update *Meta* and the graph under the iteration *i* as *G^′^*, the poisoned graph in the next iteration can be represented as:

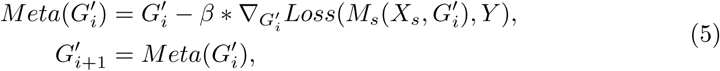

where *β* represents the learning rate, *Loss*() represents the loss function and *Y* represents the observed labels corresponding to the *X_s_* and *G*. The number of perturbation running is 10.

#### Delete internally and connect externally (DICE) attack [69]

The DICE attack randomly connects nodes with different labels or drops the edges between the nodes with the same labels, and thus we can have the perturbed graph *G^′^*. Considering the edges we intend to add are in *G^a^* and the edges we intend to preclude are in *G^p^*, this attack can be represented as:

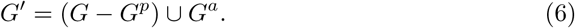

The number of perturbation running is 10.

#### Graph random attack

The graph random attack method means that we randomly add edges to the input graph. Considering the noise graph as *G^s^*, this attack can be represented as:

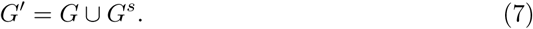

The number of perturbation running is 10.

#### Minmax attack [66]

The Minmax attack is the same as the PGD attack for a fixed GNN. This name comes from treating the perturbation as a min-max form where the inner maximization is given by the gradient ascent and the outer minimization is given by PGD, for a retrained GNN. The Minmax attack is a type of topology attack. The number of perturbation running is 10, and the attack loss type is cross entropy.

### Details of different defense methods

For the computation-oriented attack methods and biology-oriented attack methods, we consider maker gene selection and adversarial training as two methods for defending. These two approaches are applied to both *SV M_rej_* and *Cal*. For the graph-based attack methods, we consider GCNSVD, GCN-Jaccard, Robust GCN (RGCN), similarity preserving GCN (SimPGCN) for defending. These methods are ranked by the order of appearance in the main text.

#### Marker gene selection (mgs)

The mgs method, as a novel defense method proposed in this manuscript, means we train the classifier based on the clean data after selecting marker genes. Previous research [75, 93] has demonstrated that selecting marker genes can improve the model performance for annotating cell types. We use the Wilcoxon rank-sum test to discover the marker genes of each cell type, and union all the genes for training. The number of marker genes per cell type is defined in the main text. We also record the scores for exploring the contributions of different marker genes toward the cell-type classification task. If we define the clean dataset as D, the mgs approach can be represented as:

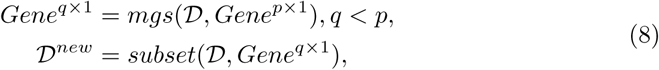

where *Gene^p×^*^1^ represents the gene list whose length is *p*, and *Gene^q×^*^1^ represents the marker gene list. *subset*() is a function to subset the expression profile by selected genes (as features). D*^new^* is used for training the classifier.

#### Adversarial training [35]

The adversarial training means that we can include the clean data and the poisoned data in the training dataset to retrain the classifier, and thus the classifier learns the information from the attack and it can perform better in the testing dataset. This defense method can be represented as:

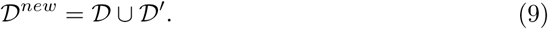

Here D*^new^* is used for training the classifier. The limitation of this method is presented in the out-of-distribution case, as it can only be used to defend the data affected by the included attacks, not for other attacks.

### Composite defense

The composite defense method means we combine the mgs method and adversarial training method together by selecting gene expression profiles with marker genes for adversarial training.

#### GCNSVD [77]

The idea of GCNSVD is to train a GCN based on the low-rank approximation of both the graph after perturbation and the feature matrix. Therefore, the model will be vaccinated and become more robust after processing.

#### GCN-Jaccard [78]

The idea of GCN-Jaccard comes from a series of observations of the behavior of the GCN model under different attacks. We first compute the Jaccard similarities in the feature space of node pairs and drop the edges existing in the nodes with many similarities. We then train a GCN with the new graph.

#### RGCN [79]

The idea of RGCN is to learn the representations of nodes using Gaussian distributions rather than vectors, and thus the model after training can automatically reflect the attacks.

#### SimPGCN [80]

The idea of SimPGCN comes from the drawbacks of the aggregation process in the GNN training, which may destroy the node similarity in the original feature space. Therefore, SimPGCN also includes a kNN graph based on the feature space and preserves this graph in the training process with an extra loss function.

### Details of evaluation methods

Here we consider both cell-type classification and spot-type classification as tasks for evaluating both attack method and defense methods. Therefore, we consider Accuracy, weighted Precision (Precision), weighted Recall (Recall), and weighted F1-score (F1-score) as metrics for evaluations. These methods are widely used in evaluating the classification problem, especially in the class imbalance case [62, 94]. The implementation of these metrics can be found in scikit-learn [62].

### Details of experimental settings

In this section, we discuss our experimental settings for evaluating both the attack methods and defense methods under different cases, as well as our methods to explore the contributions of different marker genes.

For the computation-oriented attack and biology-oriented attack, we train a classifier based on the clean dataset as a base model and adjust the clean dataset with different attack methods. We then evaluate the performances of the base model based on different poisoned datasets to reflect the robustness of the base model and the strength of different attack methods. Here the testing dataset for attacks is a poisoned dataset generated from the clean dataset. We also perform PCA to reduce the dimensions of gene expression profiles and perform the analysis for cell clustering.

For defense methods including marker gene selection and adversarial training, we first obtain the poisoned dataset from the clean dataset by running one attack method and then split the poisoned dataset into the training dataset and testing dataset (the validation dataset will be then separated from the training dataset). The test size is 0.33 for the Pancreas dataset and the PBMC dataset referred from [62, 95], while 0.2 for the Aorta dataset referred from [96]. We train the classifier based on the clean dataset and the training poisoned dataset with a defense method, and then evaluate the classifier based on the testing poisoned dataset. We repeat this process for every attack method and every defense method.

For the graph-based attack methods, we also train a classifier based on the clean graph and run attack methods for the clean graph to obtain a poisoned graph. We then evaluate the performance of this classifier based on the poisoned graph to compare the robustness of classifiers as well as the strength of different attack methods.

For the graph-based defense methods, we directly train a classifier with a defense method based on the training poisoned dataset from one attack method, and then evaluate the performance of the robust classifier based on the testing poisoned dataset. The test size is 0.33 referred from [62, 95] for both the Slide-seq v2 dataset and the Xenium dataset. We repeat this process for every attack method and every defense method.

For the analysis of marker-gene contributions, we first consider computing the overlap of marker genes across the profiles with and without attacks and then run the GO enrichment analysis to compute the enrichment ratio of all cell types. Our threshold for a significant pathway is 0.05 based on the p-value adjusted by Benjamini-Hochberg. Moreover, we consider using the single-gene attack and iteratively attacking each marker gene of different cell types from the Pancreas dataset. The accuracy after the attack for each gene is recorded to perform the comparisons.

### Data preprocessing

We follow the normalization and log1p transformation process according to Scanpy [76] to preprocess the scRNA-seq datasets and spatial transcriptomic datasets in this manuscript. For scRNA-seq datasets, We further select highly variable genes to subset the given datasets.

Although RobustCell is not biased on gender, races, and other factors, the users are solely responsible for the content they generate with models in RobustCell, and there are no mechanisms in place for addressing harmful, unfaithful, biased, and toxic content disclosure. Any modifications of the models should be released under different version numbers to keep track of the original models related to this manuscript. The users must comply with the laws of the country in which they are located.

## Data availability

We did not generate new sequencing datasets in this project. The download links and data statistics are summarized in Supplementary Data 3.

## Reproductivities and codes

To run experiments based on RobustCell, we rely on Yale High-performance Computing Center (YCRC) and utilize one NVIDIA A5000 GPU with up to 30 GB RAM for running all the attack and defense methods.

The codes of RobustCell can be found in https://github.com/HelloWorldLTY/RobustCell. The license is MIT license.

The target of current RobustCell only serves for academic research. The users cannot use it for other purposes. Finally, we are not responsible for any effects produced by other users.

## Acknowledgements

We appreciate the comments, feedback, and suggestions from Wei Jin for developing an effective attack-defence framework. We acknowledge the funding support from NIH grants P50 CA196530, U01HG013840, and R01DA065184 awarded to Prof. Zhao.

## Author contributions

T.L. designed this study. T.L., Y.X. and X.L. designed the framework. T.L. and Q.K. ran all the experiments. T.L., X.L. and H.Z. wrote the manuscript. H.Z. supervised this work.

