## Supplementary material for "RobustCell: A Model Attack-Defense Framework for Robust Transcriptomic Data Analysis": dataset: Supplementary figures.pdf

### A Supplementary figures

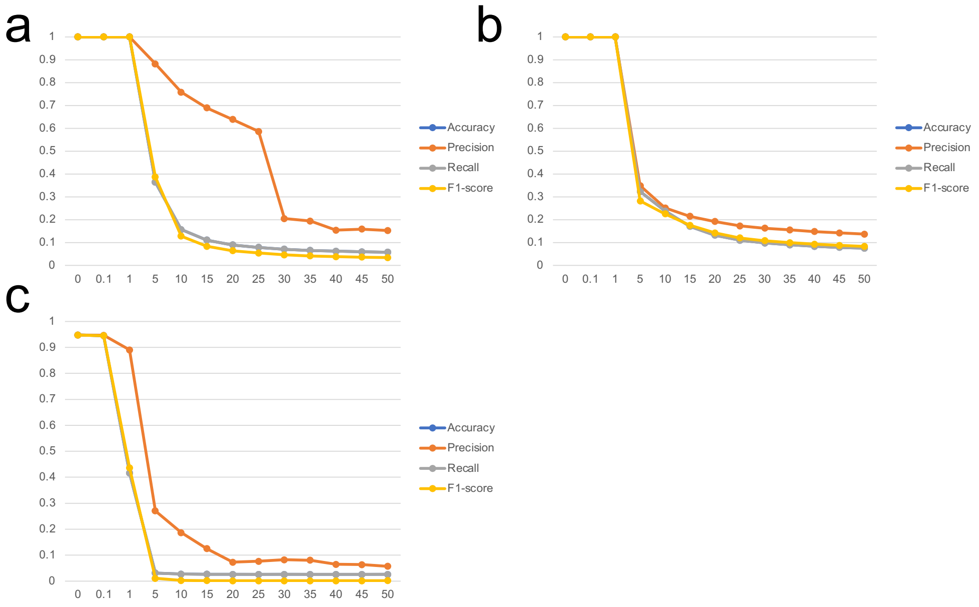


**Extended Data Fig. 1** Classification results of $SVM_{rej}$ under different types of attack methods. (a) The relationship between the attack strength *eps* of the random attack and the metrics of cell-type classification based on the Pancreas dataset. (b) The relationship between the attack strength *eps* of the random attack and the metrics of cell-type classification based on the Aorta dataset. (c) The relationship between the attack strength *eps* of the random attack and the metrics of cell-type classification based on the PBMC dataset.

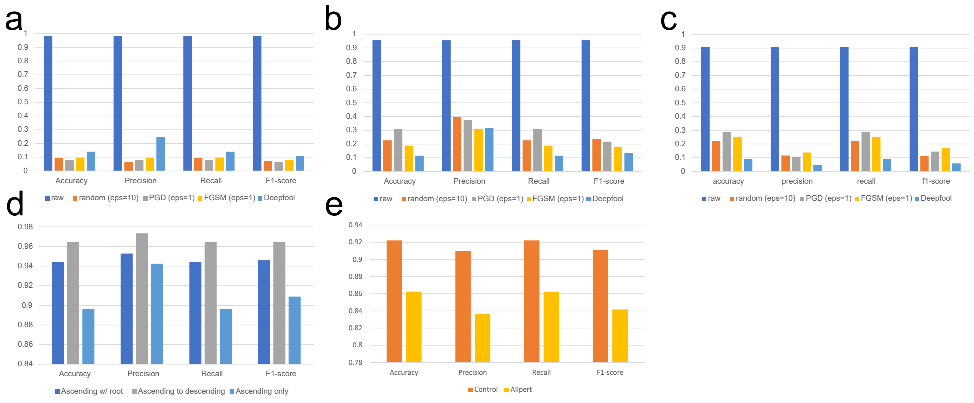


**Extended Data Fig. 2** Classification results of scGPT under different types of attacks. (a)-(c) represent the results based on computation-oriented attacks across the Pancreas dataset, the Aorta dataset, and the PBMC dataset. (d) and (e) represent the results based on biology-oriented attacks across the Aorta dataset and the Openproblems dataset.

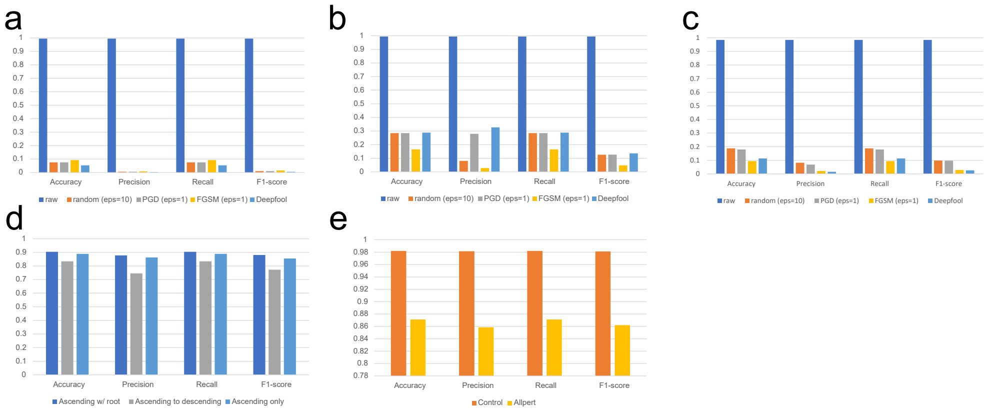


**Extended Data Fig. 3** Classification results of Geneformer under different types of attacks. (a)-(c) represent the results based on computation-oriented attacks across the Pancreas dataset, the Aorta dataset, and the PBMC dataset. (d) and (e) represent the results based on biology-oriented attacks across the Aorta dataset and the Openproblems dataset.

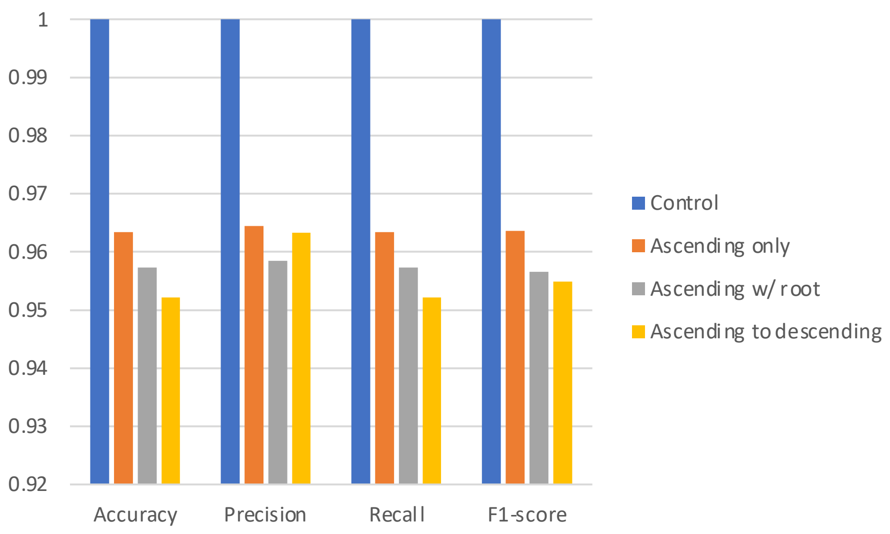


**Extended Data Fig. 4** Classification results of $SVM_{rej}$ under different types of diseases.

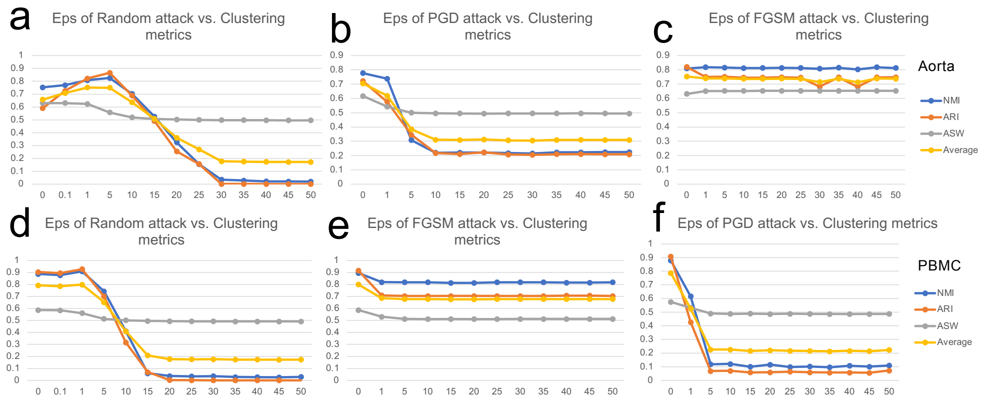


**Extended Data Fig. 5** Metrics of cell clustering under different types of attack methods for the Aorta dataset and the PBMC dataset. (a)-(c) represent the relationship between the attack strength *eps* and the metrics of cell clustering for the Aorta dataset. The attack method is mentioned in the title of each panel. (d)-(f) represent the relationship between the attack strength *eps* and the metrics of cell clustering for the PBMC dataset. The attack method is mentioned in the title of each panel.

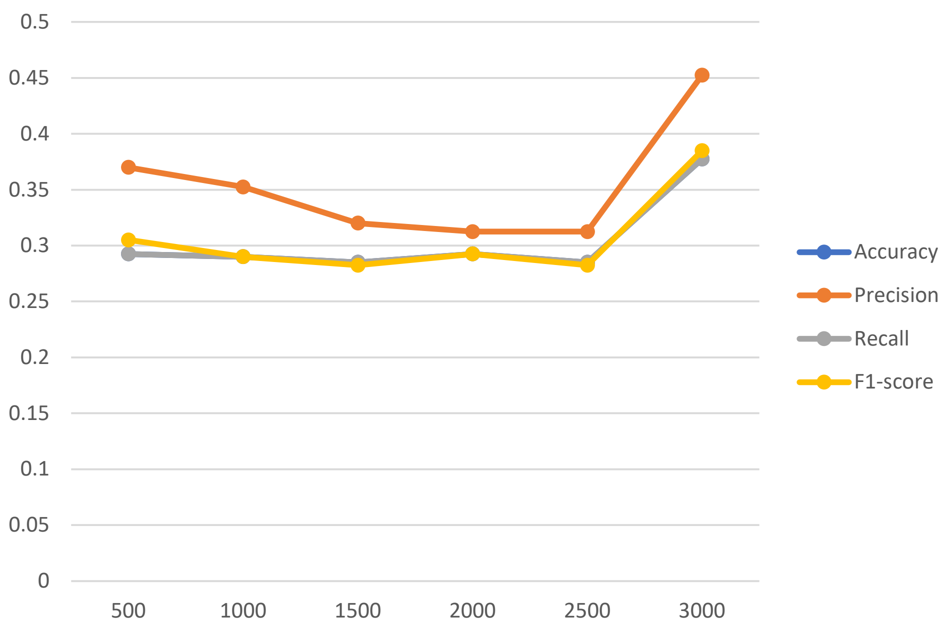


**Extended Data Fig. 6** The relationship between the number of HVGs and the performances of classification metrics averaged by different attack methods based on the Pancreas dataset.

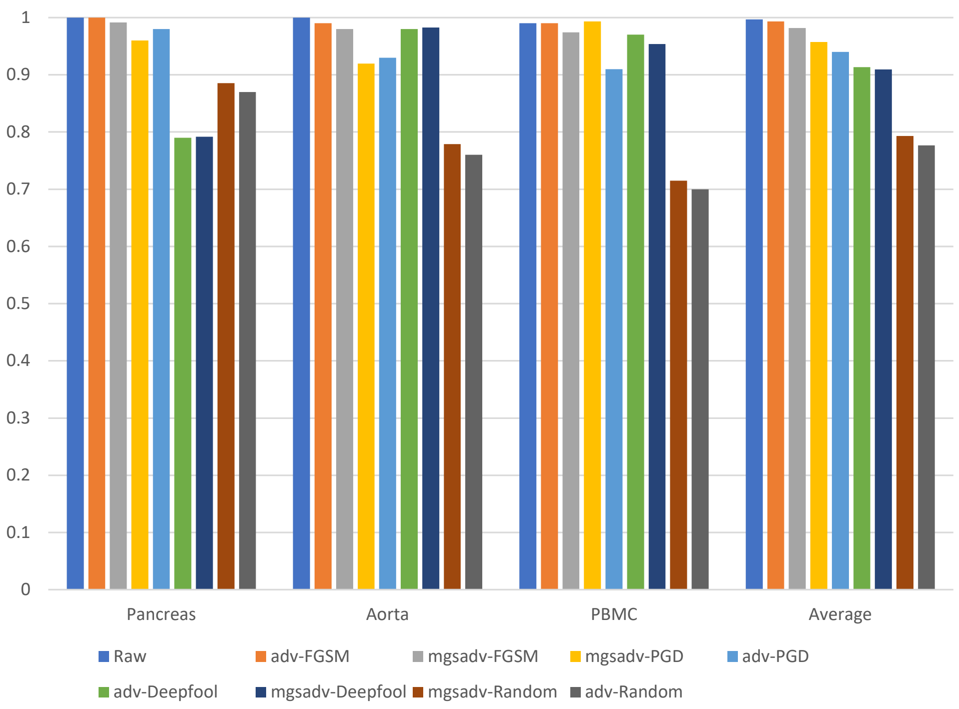


**Extended Data Fig. 7** Results of applying composite defense and adversarial training for defending different computation-oriented attack methods. We show accuracy in this figure, and the methods are ranked based on averaged accuracy across different datasets.

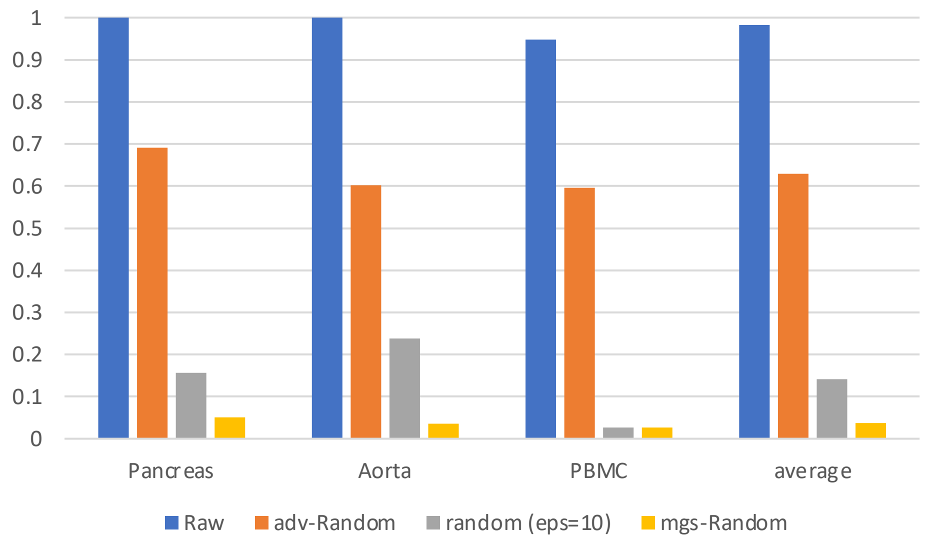


**Extended Data Fig. 8** Results of applying defense methods to $SVM_{rej}$ across different datasets. Here we display the accuracy score under different methods in the y-axis, and the bars are ranked by the averaged accuracy scores of each method.

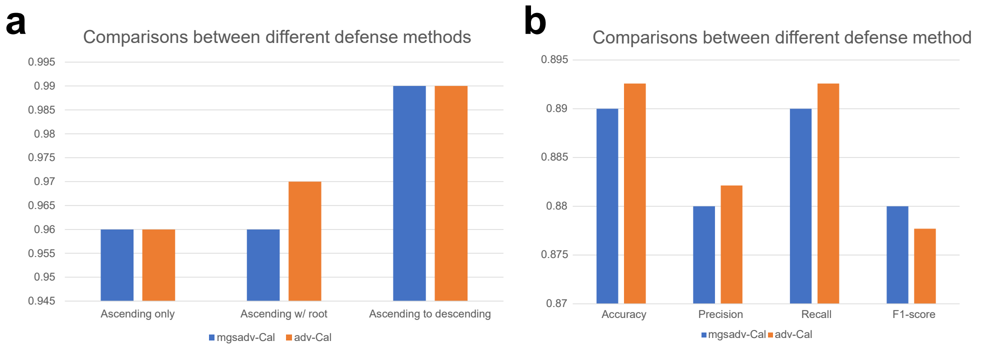


**Extended Data Fig. 9** Results of applying composite defense and adversarial training for defending different biology-oriented attack methods. (a) Comparisons between different defense methods across diseases based on the Aorta dataset by accuracy. (b) Comparisons between different defense methods for defending perturbations based on the Openproblems dataset by metrics for classification.

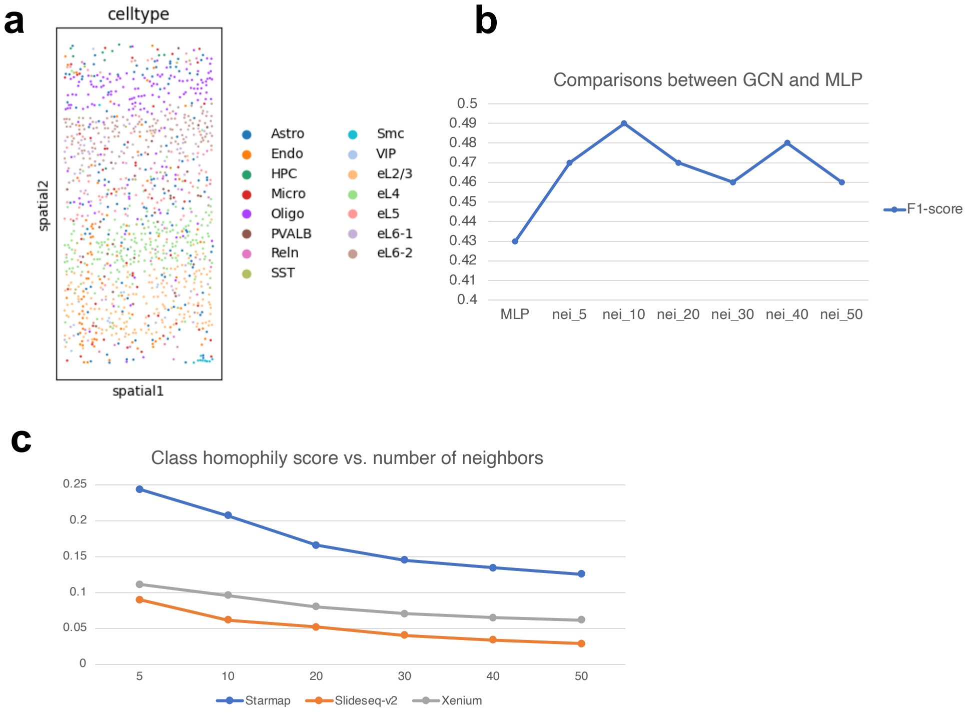


**Extended Data Fig. 10** Results of comparing GCN and MLP for spot-type classification. (a) The distribution of cell (spot) types of the Starmap dataset visualized by locations. (b) The comparison between GCN and MLP for classifying spot types by F1-score. The x axis represents the number of neighbors ($k$) used to construct the kNN graphs as well as the choices of models. (c) The relationship between class homophily scores and number of neighbors for the Slide-seq v2 dataset and the Xenium dataset.

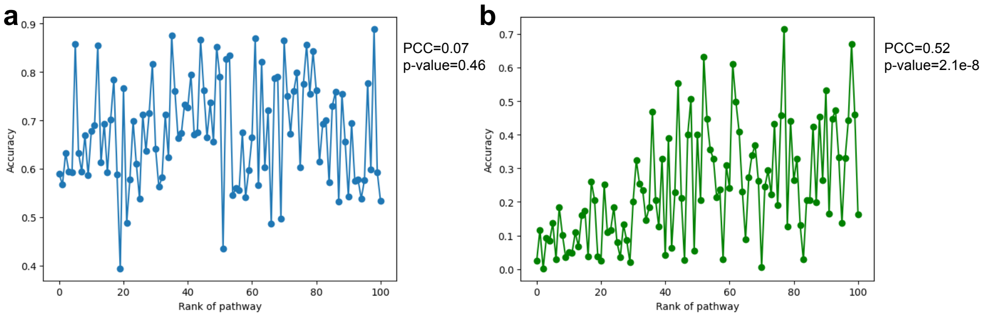


**Extended Data Fig. 11** Results of pos attack and neg attack targeting on genes in the same pathway. The pathways are ranked based on the adjusted p-values in ascending. (a) The relationship between accuracy and the rank of pathway under pos attack. (b) The relationship between accuracy and the rank of pathway under neg attack.
